# A Variational Graph Partitioning Approach to Modeling Protein Liquid-liquid Phase Separation

**DOI:** 10.1101/2024.01.20.576375

**Authors:** Gaoyuan Wang, Jonathan H Warrell, Suchen Zheng, Mark Gerstein

## Abstract

Graph Neural Network (GNN)s have emerged as a powerful general-purpose tool for representation learning across many domains. Their efficacy often depends on having an optimal underlying graph for prediction. In many cases, the most relevant information comes from specific subgraphs. In this work, we introduce a novel GNN architecture, called Graph Partitioned GNN (GP-GNN), designed to partition graphs during the prediction process, thereby focusing attention on subgraphs containing the most relevant information. Our approach jointly learns task-dependent graph partitions and representations, making it particularly effective for predictive tasks where critical features may reside within initially unidentified subgraphs. Protein Liquid-Liquid Phase Separation (LLPS) is an important physical problem well-suited to our architecture, primarily because protein sub-domains called intrinsically disordered regions (IDR)s are known to play a crucial role in the phase separation process. LLPS plays an essential role in cellular processes and is known to be associated with various diseases (*e*.*g* ., neurodegenerative diseases and cancer). However, our ability to accurately predict which proteins undergo LLPS remains limited. In this study, we demonstrate how GP-GNN can be utilized to accurately predict LLPS by partitioning protein graphs into task-relevant subgraphs, such as those highlighting IDRs. Our model not only achieves state-of-the-art accuracy in predicting LLPS for both regulator and scaffold proteins but also offers valuable biological insights, that can be used to guide further downstream investigations. Notably, upon examining subgraphs identified by the GP-GNN, we show these are consistent with annotated IDRs.

## 1 Introduction

LLPS [1, 2, 3, 4] is a phenomena underlying vital biological processes, such as the formation of membrane-less organelles in cells. Unlike membrane-bound organelles, where lipid bilayer membranes are present to encapsulate molecules spatially, LLPS describes the case where multivalent interactions between protein chains cause the formation of a distinct phase of protein condensates. Recent studies revealed evidences of the association of LLPS with the development of human diseases, such as neurodegenerative diseases and cancer [2, 5, 6, 7].

The ability of proteins to form condensates through LLPS is believed to be largely driven by the enrichment of certain binding domains in the molecule, also called IDRs [8, 9]. It is widely accepted that cation-pi, pi-pi [10], electrostatic, hydrophobic and hydrogen bonds interactions play an important role in mediating LLPS of IDRs. Yet, a quantitative understanding of LLPS driving forces is still missing. Here, we are interested in protein structure-function prediction for modeling protein LLPS. However, the large variety of distinct IDR sequence features [11, 12] and other possible unknown driving forces of LLPS makes the task difficult for conventional physical modeling approaches, but a suitable target for machine learning tools. Having an interpretable predictive tool to distinguish proteins based on their ability to phase separate will not only help us to identify phase separating proteins but also to find the key domains and properties responsible for phase separation.

In many domain areas, GNNs have emerged as a powerful tool for representation learning of systems with rich-relational data. These include social networks[13], traffic networks[14] and biological networks [15, 16, 17]. Using GNNs to study proteins, where protein structures are for instance encoded as graphs and node features carry amino acid properties, is a promising approach which has already been shown to outperform sequence-based convolutional neural networks in different tasks [18, 19, 20].

In general, the identification and choice of the optimal underlying graph for a task is crucially important for achieving high performance with GNN. For instance, it has been observed that sparsification [21] and graph structure learning [22] can enhance performance, and that such methods can avoid the problem of ‘oversmoothing’ observed in many GNN models[22]. In particular for proteins, we expect protein function relevant information to be confined within local subgraphs, supported by the fact that many protein functions are carried-out by domains. Specifically for LLPS prediction, it may be beneficial to partition the molecule graph, and remove connecting edges between task-irrelevant regions and IDRs, thereby preventing message passing from task-irrelevant regions to the IDRs. Combining the imperative to incorporate sparsification mechanisms to mitigate against oversmoothing, and the potential benefit of emphasizing distinct protein functional domains, we are motivated to propose a novel GNN architecture, a Graph-Partitioned GNN (GP-GNN), with the ability to learn advantageous graph partitions on-the-fly, and hence learn improved graph representations. In our method, prior to the graph representation learning through message passing, edges between learned subgraphs are removed (sparsification) in an adaptive fashion, to ensure that information sharing is confined within local subgraphs in the initial layers of the network.

Previous approaches to joint graph structure and representation learning in GNNs have used graph encoders [23], alternating optimization [24] and Bayesian networks [25] to perform graph structure learning. Also related to our approach are methods such as DropEdge [21] and other sparsification approaches [26], which aim to remove task-irrelevant edges, and attention-based approaches [27], which may be viewed as soft structure learning methods. Methods such as [21] learn independent measures of edge importance, and hence cannot enforce global constraints on the predicted graph structures. Further, although generative models such as [23] and [26] can in principle learn such constraints, they do not provide a mechanism for imposing such constraints in advance, and only enforce restricted forms of sparsity constraints (such as *k*-neighborhood subgraphs [26]). Likewise, attention-based methods do not enforce global constraints or sparsity directly. In contrast to the existing methods mentioned above, our approach allows desired global characteristics to be imposed onto the resulting subgraphs that emerge.

Our architecture starts with a layer of trainable meta-features, which determine the graph structure of subsequent layers; these meta-feature vectors are combined with graph-based features as input to an arbitrary clustering algorithm in order to capture important protein domains. Then, subsequent GNN layers predict phase separation ability using the combined domain characteristics.

Optimization of our model in an end-to-end manner is challenging, due to the non-differentiability of the clustering layer. A wide range of black-box optimization techniques [28] has been developed to handle objective functions which may not be globally differentiable. Promising recent approaches include score-function methods with control variates [28], variational optimization (VO) [29], and scale-space-based continuation approaches [30]. Such approaches have been applied in the context of reinforcement learning [28], graph matching [31], hyperparameter optimization [32] and differentiable deep neural network training [29], but there has been limited application of such techniques specifically to GNN architectures. In this work, we introduce an efficient VO approach, based on optimization of an smoothed objective function using a Gaussian scale-space [33]. This approach allows us to derive an enhanced form of the smoothing-based optimization algorithm [31] that makes use of local gradient information, which is important for efficiently training the graph-based layers of our model, while not assuming the objective is globally differentiable.

We first outline our novel GP-GNN architecture and our VO framework for optimization in Section 2.2 and Section 2.3 respectively. Section 3 demonstrates the effectiveness of our model in predicting protein LLPS on both regulator and scaffold proteins, in addition to analysing the properties of the graphs learned by our model and validating the identified protein subregions against external biological annotations. Section 5 concludes with a discussion, and further results are given in Supplementary Section S1, where we apply our approach to synthetic graph classification tasks with known ground truth graph structures.

Our contributions may be summarized in four key areas as follows: First, we introduce a novel GNN model adept at addressing predictive tasks, such as protein LLPS prediction, where unknown subsystems may be crucial for information extraction. Second, we present a novel metric for assessing structural-functional similarity in protein regions in Equation (3) that can be adapted to different tasks. Third, combining the above, we develop a state-of-the-art model for LLPS protein prediction, where message passing is confined within learned subregions of proteins; our model is shown to outperform all prior methods for LLPS prediction. Lastly, we corroborate the biological significance of the protein regions identified by our model by comparing them with annotated protein disordered regions and polar residue rich regions.

## 2 Methods

Our overall approach is illustrated in Figure 1 and summarized in the following. Detailed descriptions of our novel graph architecture and VO approach are given in Section 2.2 and Section 2.3 respectively.

**Figure 1:**
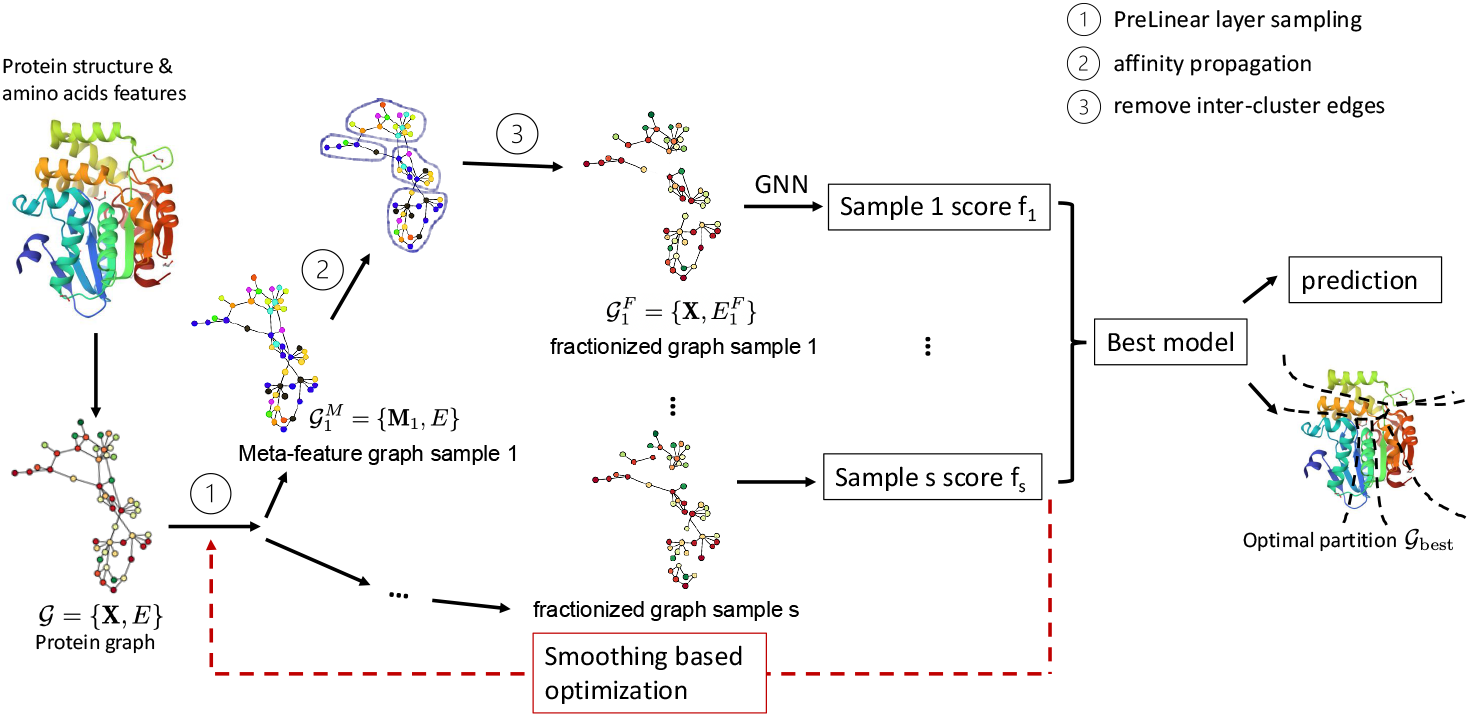
The GP-GNN architecture with adaptive cluster layers illustrated. For meta-epoch *t, N*_*S*_ samples of *Pre-Linear* layer (PLL) parameters are drawn from a multivariate normal distribution with mean *µ*_*t*_ and covariant matrix (*σ*_*t*_)^2^**I**. For each sample, we perform affinity clustering to obtain an unique set of refined graphs. We train a GNN model on the refined graphs and use the training score of the epoch with the best validation accuracy as the score function of this sample to update *µ* and *σ* using Equation (7).

1. Each element in the dataset is encoded by a graph 𝒢 = *{***X**, *E}*, where node *i* carries feature vector **x**_*i*_. For the LLPS protein prediction task, each protein is a graph 𝒢.
2. Initialize the variational distribution *Q*_*t*=0_(*θ*) at meta-epoch *t* = 0.
  2a) Draw *s* samples for the *Pre-Linear* layer’s weights, 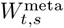, from the variational distribution *Q*_*t*_(*θ*). Applying the weights of sample *s* yields a meta-feature vector for each data-point, 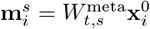, which form the rows of the meta-feature matrix, **M**_*s*_.
  2b) For each node pair *i* and *j*, a similarity score is calculated for each sample as 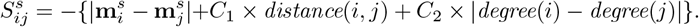 In constructing 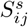, we combine a node similarity score based on the sampled meta-features **m**^*s*^, with a static graph similarity score based on closeness within the protein structure graph (including a distance and structural similarity term).
  2c) Using Affinity Propagation with the similarity matrix obtained from 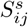, each protein graph is partitioned into subgraphs. Edges between different subgraphs are removed, resulting in a fractionized graph 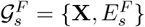
  2d) A conventional GNN is trained on 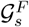 and the negative loss serves as the performance score *f*_*s*_ for each 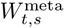 sample, corresponding to a specific graph partition.
  2e) Update the variational distribution *Q*(*θ*) via Smoothing-based optimization (SBO) based on the performance scores. Then, repeat 2a) to 2d) until convergence is attained.
3. The final prediction is made using the optimal graph partition 𝒢_best_ and the optimized GNN.

### 2.1 Protein graph generation

Our code for protein graph generation is publicly available ^1^. In our graphs, each node represents an amino acid. The node features are selected based on the nature of LLPS, supported by experimental evidence and existing physical models (see [34, 35]). We use five features: charge [36], hydrophobicity [37], polarity [38], flexibility index [39] and IDR scale [40]. The edges of the graph represent the spatial proximity of amino acids in the original molecule. In addition to the polypeptide backbone, edges are placed between pairs of nodes if: 1) their side chains exhibit specific interactions, and 2) the distance between their closest atoms is less than a threshold specific to each interaction. The considered interactions include hydrophobic, disulfide, hydrogen bond, ionic, aromatic-aromatic, aromatic-sulfur, cation-pi, and planar pi-pi interactions [10]. These interactions contribute to the molecule’s 3D conformation. All interactions are considered static, binary (unweighted), and undirected.

### 2.2 Adaptive Cluster-based Architecture

We assume that our data consists of graphs of the form 𝒢 = {**X**, *E*}, where the graph structure is given by the preliminary set of edges *E* ⊂*N* × *N*, where *N* is the set of nodes, and **X** denotes the matrix of *D*-dimensional features associated with the *N* nodes in the graph, hence for node *i* we denote its associated feature vector as **x**_*i*_ ∈ ℝ^*D*^. Additionally, for each data-point, we have an associated binary label *Y* ∈ *{*0, 1*}*.

We outline our architecture below assuming GCNConv-based message-passing [41] for convenience, although in the experimentation we also consider TransformerConv [42] message-passing formulations. Here, we let *L* denote the number of layers, while 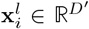 is the node feature vector of node *i* in layer *l*, and *W*^*l*^ is the layer-specific trainable weight matrix. In addition, we assume we have a further set of weights *W* ^meta^ ∈ ℝ^*D×D*^ which we use to generate a vector of ‘meta-features’ **m**_*i*_ associated with each node, and a clustering algorithm 𝒜 : ℝ ^*N×D N*^ → 𝒦^*N*^, which returns a cluster index (𝒦 = { 1…*N*_*K*_) for each node depending on the input collection of meta-features. Finally, 𝒩 (*i*, **M**) returns the neighbours of node *i* in our model, which here include only those points connected to *i* by an edge in *E* belonging to the same cluster as determined by the matrix of meta-features **M**. Our model can thus be summarized as:

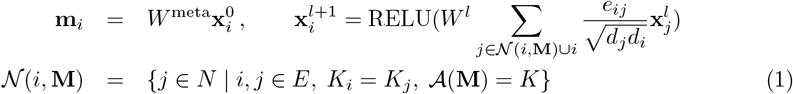

with *d*_*i*_ = 1 +Σ _*j*∈𝒩 (*i*)_, **X**^0^ = **X**, and RELU(*a*) = max(*a*, 0). We refer to *W* ^meta^ as a ‘*Pre-Linear* ‘ layer, since it is a linear transform of the input features. Also, as noted below, we apply batch correction to each layer, which simply applies a linear transformation to each *W*^*l*^ matrix. Finally, we predict the model output by pooling the final node features and multiplying by a dense output layer, *W* ^final^ ∈ ℝ^*D×*1^:

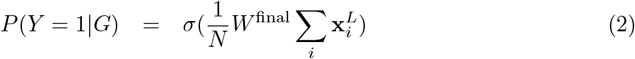

where *σ* is the sigmoid function, *σ*(*a*) = 1*/*(1 + exp(−*a*)).

### Clustering Algorithm

In this work, we use affinity propagation (AP) as our clustering algorithm 𝒜 [43]. Using meta-feature vectors **m**_*i*_ and the preliminary graph topology defined by *E*, we calculate a similarity matrix for each graph. We propose to use the following definition of similarity to capture how likely node *i* and *j* belong to the same cluster:

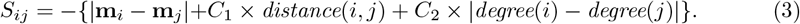

The shortest-path distance between two nodes *distance*(*i, j*) is defined as the number of edges in *E* along the shortest-path between *i* and *j. degree*(*i*) is the degree of node *i* defined as the number of edges in *E* incident this node. *C*_1_ and *C*_2_ are two scaling factors regulating the relative importance of each term. 𝒜 (**M**) is then set to the clustering returned by the AP algorithm, using similarity matrix *S* as input.

### 2.3 Smoothing-based Variational Optimization Framework

We assume we have a dataset 𝒟 consisting of pairs (*G, Y*) of matching graphs and labels. During training, our goal is thus to optimize the log-probability of our dataset: *F* (*θ*) = Σ _(*G*,*Y*)∈ *𝒟*_ log *P* (*Y* |*G, θ*), where *θ* = *{W* ^1…*L*^, *W* ^meta^, *W* ^final^*}*. Since *F* includes discontinuities due to the clustering algorithm 𝒜, we introduce a variational distribution *Q*(*θ*) = 𝒩 (*θ* | *µ, σ*^2^**I**), with **I** the identity matrix, where 𝒩 (· | *µ*, Σ) is a multivariate Gaussian distribution, and an associated smoothed variational objective:

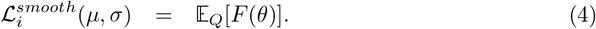

By optimizing ℒ^*smooth*^ we are optimizing a lower-bound on ℒ, since 𝔼_*Q*_[*F* (*θ*)] ≤ max_*θ*_ *F* (*θ*). Since *F* is continuous within local regions (i.e. for a fixed *W* ^meta^), we can further strengthen the smoothing objective above using local optimization. We assume we have an effective local optimizer 𝒪 for which 𝒪 (*θ*) = *θ*^*′*^ such that *F* (*θ*^*′*^) *F* (*θ*). Then we can define an augmented variational objective as:

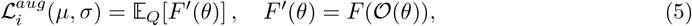

which provides a tighter bound on the original loss, since:

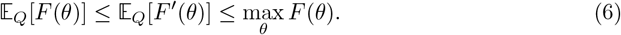

To optimize 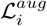, we use smoothing-based optimization (SBO) [31], combined with local gradient descent. Hence, we begin by fixing the variational distribution, {*µ*_*t*_, *σ*_*t*_} at *t* = 0, and draw samples 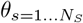 from 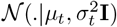 We then apply our local gradient descent optimizer 𝒪 to each sample, which fixes the meta-features while optimizing the subsequent layers of the GNN, generating 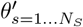 We calculate 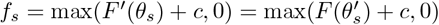

#### Algorithm 1

Smoothing-based Optimization Augmented with Local Gradients

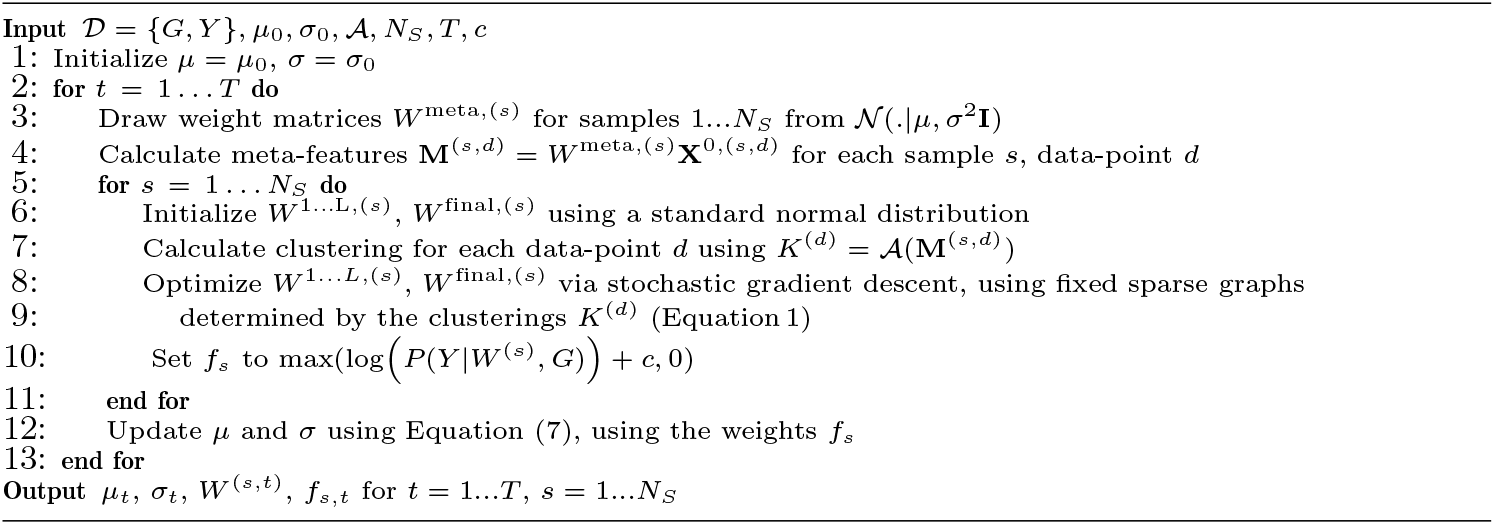

where *c* is an offset to ensure that *f*_*s*_ is non-negative, and calculate *µ*_*t*+1_ and *σ*_*t*+1_ using the following updates:

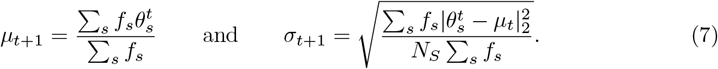

We note that, although the updates in Equation (7) may be applied simultaneously as in [31], we found that applying them sequentially improved the convergence of the algorithm to a local optimum. Also, in practice we use the default PyTorch-Geometric parameter initialization settings for *W* ^1…L^ and *W* ^final^, and restrict the distribution *Q* to the parameters *W* ^meta^; this is equivalent to setting *µ*_*i*_ = 0 for all other parameters, and using two separate standard deviations, *σ*_*a*_ and *σ*_*b*_ for *W* ^meta^ and all other parameters respectively, where only *σ*_*a*_ is updated according to Equation (7), and the second remains fixed at *σ*_*b*_ = 1 (hence all other parameters are always initialized using a standard normal distribution). We summarize our smoothing-based local stochastic gradient descent algorithm in Algorithm 1.

## 3 Results

We apply our model to two LLPS tasks, involving predicting protein LLPS for regulator and scaffold proteins. In addition, Supplementary Section S1 gives further results on a synthetic graph classification dataset designed to test the assumptions underlying our model. The synthetic dataset is designed to provide a classification task similar to the protein LLPS task, but with known ground-truth functional domains. It preserves identical overall statistical characteristics across both positive and negative instances; however, the distribution of the features is aligned with the graph clusters only in the positive graphs. Using this synthetic dataset, we further evaluate our model’s ability to detect local features within subregions of the graph. We first comment on the common features of the architectures tested (including implementation and compute time), before outlining the specific experimental setup and results for each task.

### Datasets

LLPS occurs when multivalent interactions between protein chains cause the formation of a distinct phase of protein condensates. The ability of proteins to form condensates through LLPS is believed to be largely driven by the enrichment of certain binding domains in the molecule, such as the intrinsically disordered regions (IDRs) [8]. IDRs [9] are considered to be segments that lack unique 3D structures. We prepare three LLPS datasets focusing on regulator and scaffold proteins. Basic protein statistics for the three datasets, including chain length and amino acid composition, can be found in Supplementary Section S2.1.

The first dataset is the *Regulator PDB* dataset, where the input protein graphs are constructed using ground truth PDB 3D structures. For positive examples in this regulator dataset, we use the 135 proteins labeled as regulators in the DrLLPS dataset [44]. We kept only those proteins for which the spatial coordinate information of at least 70% of the chain is present in the respective PDB file. If multiple PDB files are linked to one protein, we choose the one with the highest spatial resolution. Our dataset contains an equal number of negative examples, which are selected randomly from Uniprot [45] such that they are not found in the DrLLPS dataset, and their PDB files are required to contain the full chain spatial information.

The second dataset is the *Regulator AlphaFold* dataset, where the AlphaFold predicted 3D structures are used to construct the protein graphs. This dataset contains nearly identical

sets of regulator proteins as the previous one, while the latter uses the ground truth PDB 3D structure to construct the protein graphs. The third dataset is the *Scaffold AlphaFold* dataset, containing 137 proteins labeled as scaffold in the DrLLPS dataset [44], for which

we use the AlphaFold predicted 3D structures to construct the protein graphs. We then select an equal number of negative samples that are not found in the DrLLPS dataset and have AlphaFold predicted 3D structure information on UniProt webpage. See supplementary material for full list of proteins used. We divide the data into 40% training, 30% validation and 30% test set.

### Model Architectures and training

In this work, we consider several realizations of the GP-GNN framework. The *Pre-Linear* +Clustering+GNN (PCG) models represent full implementations of the proposed GP-GNN framework, while the Clustering + GNN (CG) models are partial implementations that do not include the adaptive PLL component. We further divide the two classes into subcategories based on the message passing operator used, including GCNConv [41] and TransformerConv [42] based models, and name them based on the combination of the class and the message passing operator. For GCNConv-based models, the subcategories were named CGCN, and PCGCN. For TransformerConv-based models, the subcategories were named CGTN, and PCGTN. In addition, we include conventional GNN architectures as baseline models, referred to as basic GCN and GTN models.

We test all graph-based methods on the same datasets, ensuring they have equal access to the same information. This prevents our model from gaining any advantage due to potential biases between the positive and background datasets. Additionally, we develop a logistic regression model using only basic protein statistics, including chain length and amino acid composition, as a baseline. The basic GNN models and the GNN module embedded in CG and PCG models have four layers. They use the Adam optimizer with a learning rate of 0.008, a batch size of 30, and the dimension of the node features of all hidden layers is set to 25. The maximal number of epochs is set to 300 and the validation accuracy is monitored with a patience parameter of 160 to determine early stopping. This set of hyperparameters was found to be optimal for the stand alone basic GCN model on LLPS dataset. For the CG and PCG models, we use AP with a pre-defined similarity matrix to perform the clustering. The off-diagonals of the similarity matrix are calculated with Equation (3). The diagonals are filled with *S*_*m*_, the median of all *S*_*ij*_, for *i* ≠ *j*. The damping factor of the AP is set to 0.88 and the maximum number of iterators is set to 1600. We set the preference parameter of the AP to *p*_fac_ *× S*_*m*_. We test various combinations for *cp* = (*p*_fac_, *C*_1_,*C*_2_). Recall that *C*_1_ and *C*_2_ are factors from Equation (3) controlling the relative importance of different terms. We noticed that increasing *p*_fac_ will result in larger average cluster sizes and increasing *C*_1_ leads to higher node degree.

For all PCG models, SBO is used for the PLL training as described in Section 2.3. In each meta-epoch, the number of random samples is set to *N*_*S*_ = 15. The score function used to update *µ* and *σ* is derived from the CrossEntropyLoss of the training dataset *L*_CE_(train) according to max(− *L*_CE_(train) + 0.6, 0). In our implementation, each meta-epoch consists of an update of *µ* followed by an update of *σ*.

For basic GCN/GTN and random dropout GCN/GTN, we report the average accuracy over all samples as the performance. For CG and PCG models, we report the test accuracy of the model with the best validation accuracy across all meta-epochs, as the top model performance. The average test accuracy of the meta-epoch with the highest average validation accuracy is reported as the average performance. To allow an exact comparison with the PCG models, we separate the results of the CG models also into groups, each consisting of 15 samples.

We repeat the training of basic GNN models with 105 random initial parameters. For the CG models, the training is repeated 105 times and separated into 7 groups in analogy to the meta-epochs. For PCG models, we train the models for 60 meta-epochs. We initialize *µ* such that the PLL is equal to identity (equivalent to CG) and 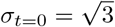. At the end of the training, we observe that the final *σ* around *t* = 60 is typically well below 0.1. During our hyper-parameter grid search for *cp*-parameters, we covered a range of cluster sizes from less than 10 to over 60 nodes and approximately 10% to 63% of the full graph node degree for regulator proteins. For the scaffold proteins, we tested a much larger range of *cp*-parameters and covered cluster sizes from ≈10 to ≈70 and ≈20% to ≈80% of the original graph node degree.

### Implementation and compute time

Open-source python libraries including Torch, PyTorch-geometric and NetworkX are used in the implementation. The message passing operators TransformerConv and GCNConv from torch geometric.nn are used. The AP is performed using the function sklearn.cluster.AffinityPropagation. For protein graph generation, we use the python package *proteingraph*[46]. With respect to compute time, one meta-epoch for one sample takes ∼ 3 hours on a CPU with 32G. For the PCG model, 60 meta-epochs for one *cp* with *N*_*S*_ = 15 required approximately 2500 CPU-hours. For the CG models, ∼ 400 CPU-hours were required for one *cp* setting.

### 3.1 Model performance on regulator proteins

In this section, we apply our model for the classification of regulator proteins. We consider two regulator protein datasets: the *Regulator PDB* dataset and the *Regulator AlphaFold* dataset described above. Additional performance tables can be found in the Supplementary Section S2.2.

#### Performance on Regulator PDB dataset

In Table 1, we summarize the performance of basic GCN/GTN models and random dropout models as the baseline performance of classification of regulator proteins using graphs constructed with (noisy) ground truth PDB 3D structures. For this dataset, the range of *cp* = (*p*_fac_, *C*_1_, *C*_2_) settings explored is: *p*_fac_ ∈ {1, 2, 3, 4, 5, 6 }, *C*_1_ ∈ {0.1, 0.2, 0.4, 0.8} and *C*_2_ ∈ {0.1, 0.4 }. In Table 2, we show the best three choices of *cp*-parameters for PCG models based on their performances on the validation dataset. PCGCN achieved a better performance than PCGTN on the validation dataset. For the best *cp*-parameters for the PCGCN, we also tested the CGCN models, namely using raw features as input for clustering instead of meta-features. Since the difference between the validation performance of these models is within the error range and the *cp*-values are very close to each other, we report the average test AUC of the top three models as the final test performance. We found that PCGCN using an adaptive cluster-based architecture with an average test AUC of 0.749 outperforms CGCN models using a fixed sparsification strategy, as well as basic GCN/GTN models without sparsification and GCN/GTN models with random sparsification through dropout.

**Table 1:** *Regulator PDB* dataset, GCN and GTN models. The test AUC is shown for both basic GCN and GTN, as well as for random dropout GCN and GTN. See also Table S2.

| drop % | basic (0%) | 50% | 70% | 90% | logistic regression |
| --- | --- | --- | --- | --- | --- |
| GCN | $0.708 \pm 0.036$ | $0.718 \pm 0.047$ | $0.717 \pm 0.058$ | $0.729 \pm 0.058$ | 0.543 |
| GTN | $0.721 \pm 0.037$ | $0.709 \pm 0.041$ | $0.697 \pm 0.043$ | $0.69 \pm 0.058$ | |

**Table 2:** *Regulator PDB* dataset, PCGCN and PCGTN models. Average performance of top 3 *cp*-parameters for PCGCN and PCGTN is presented. ‘meta/raw’ node degree (cluster size) denotes the average node degree (cluster size) obtained by using the optimized meta-features or raw features as input for the clustering algorithm. See also Table S3.

|  | test AUC | meta/raw node degree | meta/raw cluster size | CG test AUC |
| --- | --- | --- | --- | --- |
| **PCGCN** |  |  |  |  |
| 2, 0.1, 0.2 | $0.715 \pm 0.042$ | 0.551/0.615 | 30.1/ 23 | $0.707 \pm 0.052$ |
| 2, 0.1, 0.1 | $0.775 \pm 0.025$ | 0.67/0.667 | 42.5/24.9 | $0.721 \pm 0.045$ |
| 1, 0.2, 0.1 | $0.757 \pm 0.034$ | 0.593/1.101 | 19.2 / 8.19 | $0.741 \pm 0.054$ |
| **PCGTN** |  |  |  |  |
| 1, 0.1, 0.1 | $0.718 \pm 0.029$ | 0.466/0.572 | 27.36/13.13 | |
| 1, 0.2, 0.1 | $0.712 \pm 0.033$ | 0.524/1.101 | 19.2 / 8.19 | |
| 1, 0.1, 0.4 | $0.733 \pm 0.039$ | 0.362/0.382 | 20.3/15 | |

#### Performance on Regulator AlphaFold dataset

The performance of our model on the regulator dataset but using AlphaFold predicted 3D structure is presented in Table 3. We performed a grid search for *cp*-parameters with *p*_fac_∈ [7, 9], *C*_1_ ∈ [2, 4] and *C*_2_ = 0.1, and we also tested the nine most promising *cp*-values from the scaffold AlphaFold dataset (see below Section 3.2). Similar to our calculation of the performance of our model on regulator proteins with ground truth 3D structure information, we average over the three models with the best performance on the validation dataset. PCGTN demonstrated the best performance, achieving a test AUC of 0.786. One should note that the best *cp*-parameters are among the most promising ones on the scaffold dataset.

**Table 3:**
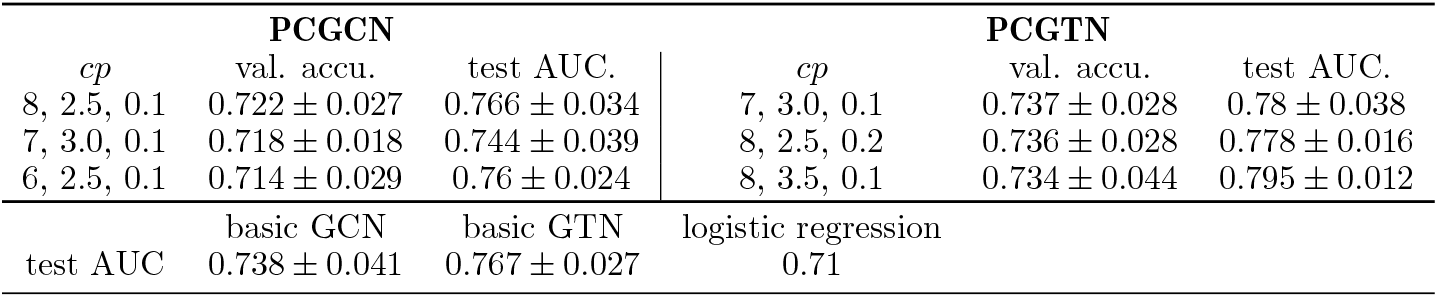
*Regulator AlphaFold* dataset, all models. Average performance of the three *cp*-parameters with the highest validation performance for PCGCN and PCGTN is presented. See also Table S4.

These results may reflect the interplay between the quality / noise characteristics of the data, and the complexity of the model. For instance, the AlphaFold predicted structures are less noisy than the PDB structures (although potentially with biases introduced by AlphaFold), which may plausibly allow the PCGTN models to achieve better performance since they are less prone to over-fitting in this context than the noisy PDB dataset. The latter, however, may favor the PCGCN, which contains fewer parameters.

In addition, we show the average validation and test accuracy as a function of meta-epochs *t* for one PCGCN and one PCGTN model for the two datasets using different protein graphs in Figure 2, where the basic GCN and basic GTN performances are also shown as baselines. The validation accuracies of our adaptive models shows a clear upwards trend especially towards the later stages of the training, suggesting that the PLL, and thus meta-features, are successfully trained through our variational optimization. The gaps between the validation accuracies and test accuracies in Figure 2 (A) indicate overfitting. This is not observed on the *Regulator AlphaFold* dataset. As discussed above, this may reflect the advantages of the more flexible PCGTN architecture on the less noisy AlphaFold structures.

**Figure 2:**
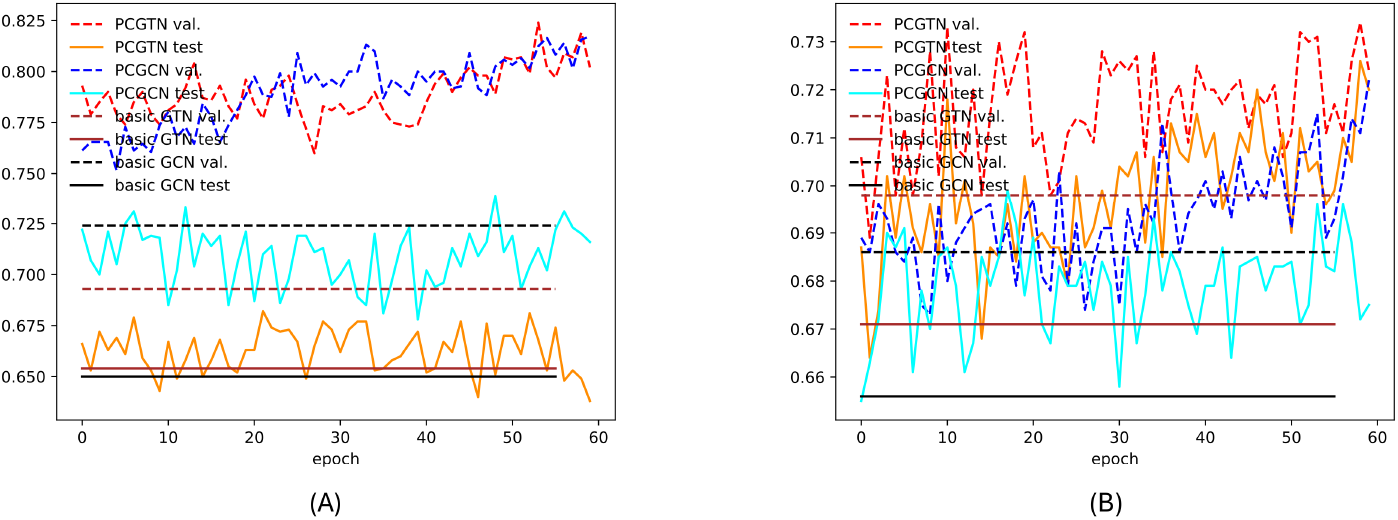
Regulator dataset, accuracy during training. Note that for basic GCN and GTN models, we show a simple average over all samples. (A): *Regulator PDB* dataset. The averaged validation (dashed) and test (solid) accuracy are shown for PCGCN *cp*=(2,0.1,0.1) and PCGTN *cp* = (1, 0.1, 0.1). The best meta-epoch of PCGTN is 53 and the best meta-epoch of PCGCN is 59. (B): *Regulator AlphaFold* dataset. For PCGCN, *cp* = (8, 2.5, 0.1) is shown with the best epoch at 59. For PCGTN, *cp* = (8, 3.5, 0.1), the best epoch is found at epoch 58.

### 3.2 Model performance on scaffold proteins

In this section, we present the performance of our model on scaffold proteins using AlphaFold-predicted 3D structures, referred to as the *Scaffold AlphaFold* dataset. A similar test on scaffold proteins using ground truth PDB 3D structure was not possible due to the very limited number of known scaffold proteins for which the ground truth 3D structure is available. To find the optimal *cp*-parameters (*p*_fac_, *C*_1_, *C*_2_), we first performed a comprehensive hyper-parameter screening for *p*_fac_ ∈ [2, 10], *C*_1_ ∈ [0.1, 20] while keeping *C*_2_ = 0.1. Then, for several *p*_fac_ and *C*_1_ choices with the highest validation accuracies, we screen for *C*_2_ over a range of *C*_2_ ∈ [0.1, 3]. The performance of our model and baseline models are listed in Table 4.

**Table 4:** Test AUC on the *Scaffold AlphaFold* dataset for various models. See also Table S5.

| basic GCN | basic GTN | random dropout GTN | logistic regression |
| --- | --- | --- | --- |
| $0.804 \pm 0.05$ | $0.856 \pm 0.047$ | $0.851 \pm 0.048$ | 0.729 |
| PCGCN(7,3.0,0.1) | CGTN(8,2.5,0.2) | PCGTN(8,2.5,0.2) average | PCGTN(8,2.5,0.2) peak |
| $0.832 \pm 0.056$ | $0.773 \pm 0.079$ | $0.894 \pm 0.033$ | 0.92 |

Additional performance tables can be found in Supplementary Sections S2.2 and S2.3.

In Figure 3, the validation accuracies resulting from our parameter screening for PCGTN *p*_fac_ and *C*_1_ is shown. The best region for both the average performance and peak performance is identified to be *p*_fac_ ∈ [5, 8] and *C*_1_ ∈ [1.0, 3.0]. For the top performing *p*_fac_ and *C*_1_ values in this region, we continued to perform a screening over *C*_2_ ∈ [0.1, 3]. Among the top performing models, we found that the difference between the validation accuracies among the top 20 models is less than 2%, which is less than the error range. We identify the small region centered around *cp* = (8, 2.5, 0.2) ^2^ to be the best *cp*-parameter choice. For *cp* = (8, 2.5, 0.2), our model achieved an average test AUC of 0.894 *±* 0.033 in the best performing meta-epoch and a peak test AUC of 0.92, outperforming basic GTN, random dropout GTN, and CGTN. ROC curves of the PCGTN model with the peak performance and the best-performing meta-epoch (calculated by combining the predictions of all models from this meta-epoch) are shown in Figure 3d.

**Figure 3:**
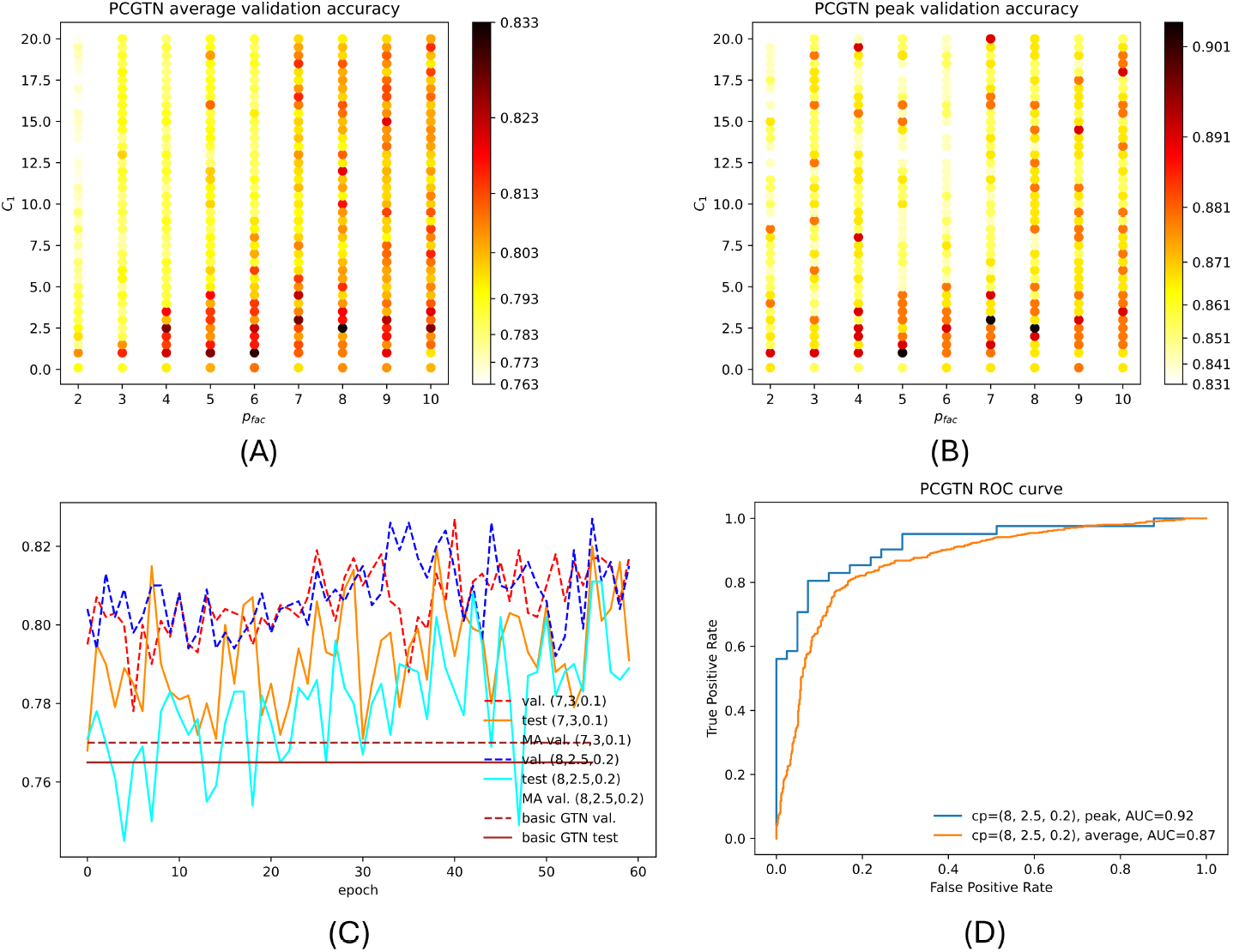
*Scaffold AlphaFold* dataset, PCGTN performance. 2D plots showing *p*_fac_ and *C*_1_ dependency of the validation accuracy of the PCGTN at *C*_2_ = 0.1; the optimal *C*_2_ value is only screened using the most promising *p*_fac_ and *C*_1_ choices. (A) average validation accuracy in the best performing epoch. (B) Peak validation accuracy. (C) Average validation accuracy and test accuracy against number of meta-epochs for the best model *cp* = (8, 2.5, 0.2) and the second best model *cp* = (7, 3.0, 0.1) compared with the performance of the basic GTN and PCGTN models, show clear improvement of the PCGTN through the variational optimization and gradual outperformance with respect to the basic GTN. The best meta-epoch of *cp* = (8, 2.5, 0.2) in this example is found to be 55 and for *cp* = (7, 3.0, 0.1) it is meta-epoch 40. (D) ROC curves for the peak performing model and the average over all models in the best metaepoch.

In addition to the PCGTN model, we provide here also the results of the PCGCN models. As shown in Figure 4. The best *cp*-parameters for the PCGCN are found to be in the region *p*_fac_ ∈ [5, 7] and *C*_1_ ∈ [0, 5]. Through additional *C*_2_ screening in this region, we found the best *cp*-parameter to be (7, 3.0, 0.1) with a test AUC of 0.832 *±* 0.056.

**Figure 4:**
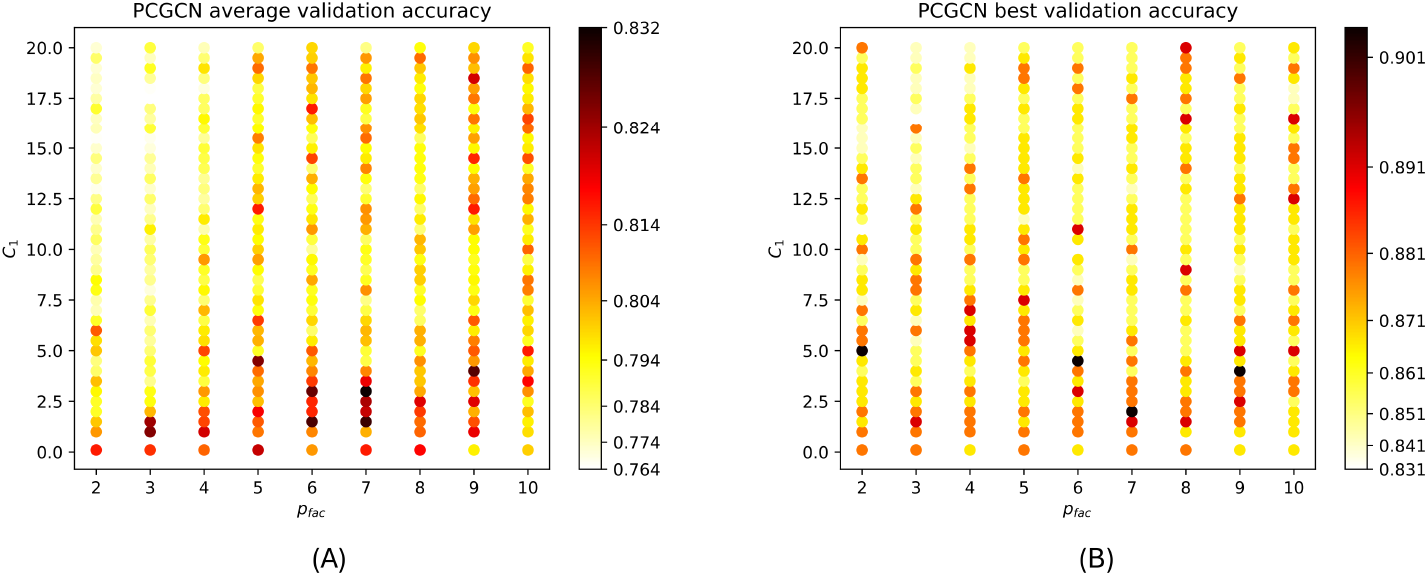
*Scaffold AlphaFold* dataset, PCGCN performance. 2D plot showing *p*_fac_ and *C*_1_ dependency of the validation accuracy of the PCGCN at *C*_2_ = 0.1; the optimal *C*_2_ value is only screened using the best *p*_fac_ and *C*_1_ choices. (A): average validation accuracy in the best performing epoch. (B): Peak validation accuracy.

We observed that the average performance of the top 20 models for PCGTN is slightly higher that of PCGCN. We also note that for the AlphaFold predicted structures, the more flexible transformer-based model (see Figure 3c) no longer has the over-fitting problem caused by the noisy nature of the ground truth PDB graph as in Section 3.1. Combining these considerations, we conclude that PCGTN achieves the best overall performance in the region around *cp* = (8.0, 2.5, 0.2), agreeing with the results on the *Regulator AlphaFold* dataset above, which showed the PCGTN to perform better than PCGCN when using AlphaFold as opposed to PDB structures, since the PCGTN, with its increased model complexity, can exploit the less noisy datasets generated by AlphaFold.

### 3.3 Comparison to previous predictors

Previous methods for phase separation protein (PSP) prediction [10, 47, 48, 49] have concentrated on predicting scaffold proteins. To our knowledge, the optimal performance using previous methods is a recall of 10.54% on regulators and 46% on scaffolds, reported in Ref. [48] at an acceptance rate of 0.02 for the background class (here, the whole proteome). We thus applied our optimal models and ranked each protein in our test set by the final score 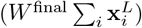 We taken into account the results of all 15 samples in the epoch of the best average validation accuracy, and found the recall at a 0.02 acceptance rate to be 23.6*±*6.8% for the PCGTN model with *cp* = (8, 3.5, 0.1) trained with the *Regulator AlphaFold* dataset and 50.6 *±* 16.4% for scaffolds using PCGTN with *cp* = (8, 2.5, 0.2), suggesting that our model achieves better performance than prior methods for the prediction of regulators and scaffolds. We further bench-marked our performance on the scaffold dataset by 1) applying three existing predictors to our dataset and 2) comparing our results with those of eleven predictors on a comparable dataset. In both comparisons, our model demonstrated performance that was either comparable to or better than the previous predictors. See Supplementary Section S2.4 for further details of the comparisons with recent sequence-based LLPS predictors.

### 3.4 Properties of the learned clusters

Since we observed above that the clustered model performed best overall, we were interested in the overall properties of the subgraphs discovered by these models.

The three best PCGCN models on the *Regulator PDB* dataset with optimized PLL have a singleton percentage of 65.12%, 59.92% and 63.72%. Although the average overall node degree is only around 0.6, the average node degree per cluster after singleton removal is 2.13, 2.45 and 2.13. For the optimized PCGTN model *cp* = (8, 3.5, 0.1) on the *Regulator AlphaFold* dataset the singleton percentage is 70.4%. The average node degree per cluster after singleton removal is 1.82. The singletons, defined as nodes without edges, can be interpreted as potentially noisy nodes, thus the model may have learned to prohibit the information propagation to their neighbors. It is worth mentioning that the optimal *cp*-combinations on the *Regulator AlphaFold* dataset are different from the *Regulator PDB* dataset. This is not surprising since the AlphaFold predicted structures only include the chain of interest, while PDB graphs often include unwanted information from other chains in the complex, or multiple conformations of the same chain.

We noticed that the cluster statistics of the best models on the *Regulator AlphaFold* dataset are very similar to those on the *Regulator PDB* dataset despite the distinct *cp*-parameters. We believe that the stability of the cluster statistics in these tests using quite different graphs reflects underlying aspects of the LLPS regulator protein biology. We note that this range of node degree is consistent with the nature of the lack of stable structure in IDRs, which is a result of missing bonds that are responsible for maintaining the 3D structure of proteins in shape. In case of a linear chain with only backbones, the average node degree is 2, which is close to the values observed in the best models above.

For the *Scaffold AlphaFold* dataset, the best performing model (PCGTN with *cp* = (8.0, 2.5, 0.2)) measured an average node degree of 2.1 and an average node degree per cluster after singleton removal of 4.8 (the singleton percentage is 24.76%).

We noticed that the best models, despite having different *cp*-values, underlying structure information and message-passing schemes, demonstrate similar cluster statistics on refined graphs through the adaption of the PLL parameters (see Table 2, which compares raw node degree and raw cluster size to meta node degree and meta cluster size). Further, we note that our comprehensive grid search of *cp*-parameters in the scaffold results shows a gradual change in performance, and reveals clearly the presence of a stable optimal region of the clustering parameters leading to enhanced classification accuracies, suggesting the above statistics are stable for the optimal models.

## 4 Validation against annotated protein regions

In this section, we assess the biological relevance of the clusters identified by our model, by comparing them against annotations from UniProt, a comprehensive resources for protein sequence and annotation data. Specifically, we analyze the performance of the PCGTN model on the *Scaffold AlphaFold* dataset, demonstrating its proficiency in identifying biological significant features crucial to protein functionality. UniProt provides several vital annotations, notably the identification of disordered protein regions. These regions, lacking a stable three-dimensional structure under physiological conditions, play significant roles in protein function despite their unordered nature.

For each protein, the model identifies *n* clusters, while UniProt provides *m* disordered regions. The overlap score and cluster size error are calculated as follows (refer to Supplementary Section S3.2 for alternative methods):

### Overlap assessment using Jaccard Index

- **Procedure**: For each disordered region from UniProt, we calculate the Jaccard Index with every cluster identified by the PCGTN model. The Jaccard Index, a measure of set similarity, is computed as the size of the intersection divided by the size of the union of the disordered region and the cluster.
- **Selection of maximum indices**: The highest Jaccard Index is selected for each disordered region, representing the best match with the model’s clusters.
- **Weighted average calculation**: We compile *m* maximum Jaccard Indices, each corresponding to a disordered region from UniProt. The final overlap score is the weighted average of these indices. Weights are proportional to the length of the disordered regions, giving greater importance to larger regions.

### Cluster size ratio error analysis

- **Error calculation for size compatibility:** We assess the size compatibility between the model’s clusters and the disordered regions by calculating the cluster size ratio error. This error is the absolute value of the difference between 1 and the cluster size ratio, defined as the size of the cluster divided by the size of the disordered region.
- **Selection and weighting of errors:** The smallest error for each disordered region is selected. The final cluster size ratio error score is the weighted average of these smallest errors, with weights based on the length of the disordered regions.

We compared the overlap score distributions between proteins in the signal (LLPS) group (labeled as scaffold in the DrLLPS dataset) and the background group (not present in the DrLLPS dataset). We divided the score range [0,1] into 50 bins, analyzing the relative frequency of scores within each bin.

Our biological interpretation considered a subset of parameters, *p*_fac_ ∈ [2, 10] and *C*_1_ ∈ [0.1, 10] (half the range of the parameter screening we did for the performance test) while keeping *C*_2_ = 0.1, then an additional screening for *C*_2_ − in the range *C*_2_ ∈ [0.1, 3] for the top-performing parameters as described in Section 3.2.

Various metrics for different *p*_fac_ and *C*_1_, and *C*_2_ = 0.1 are shown in Figure 5: the median difference between in overlap score between signal and background proteins in Figure 5(a), right-hand Mann-Whitney p-value for the hypothesis that signal proteins have higher overlap scores than background proteins in Figure 5(b), and the mean difference in the distribution of the absolute value of 1 minus the cluster size ratio, indicating optimal size match Figure 5(c). Examples of overlap score distributions for three *cp*−parameters with the best performance is shown in Figure 5(d). We note that for *C*_2_ ≠ 0.1, significant p-values and positive median differences between signal and background proteins are observed from the overlap score of the cp-parameters that correspond to the best performing models. Refer to Supplementary Section S3.1 for further results regarding *C*_2_≠ 0.1.

**Figure 5:**
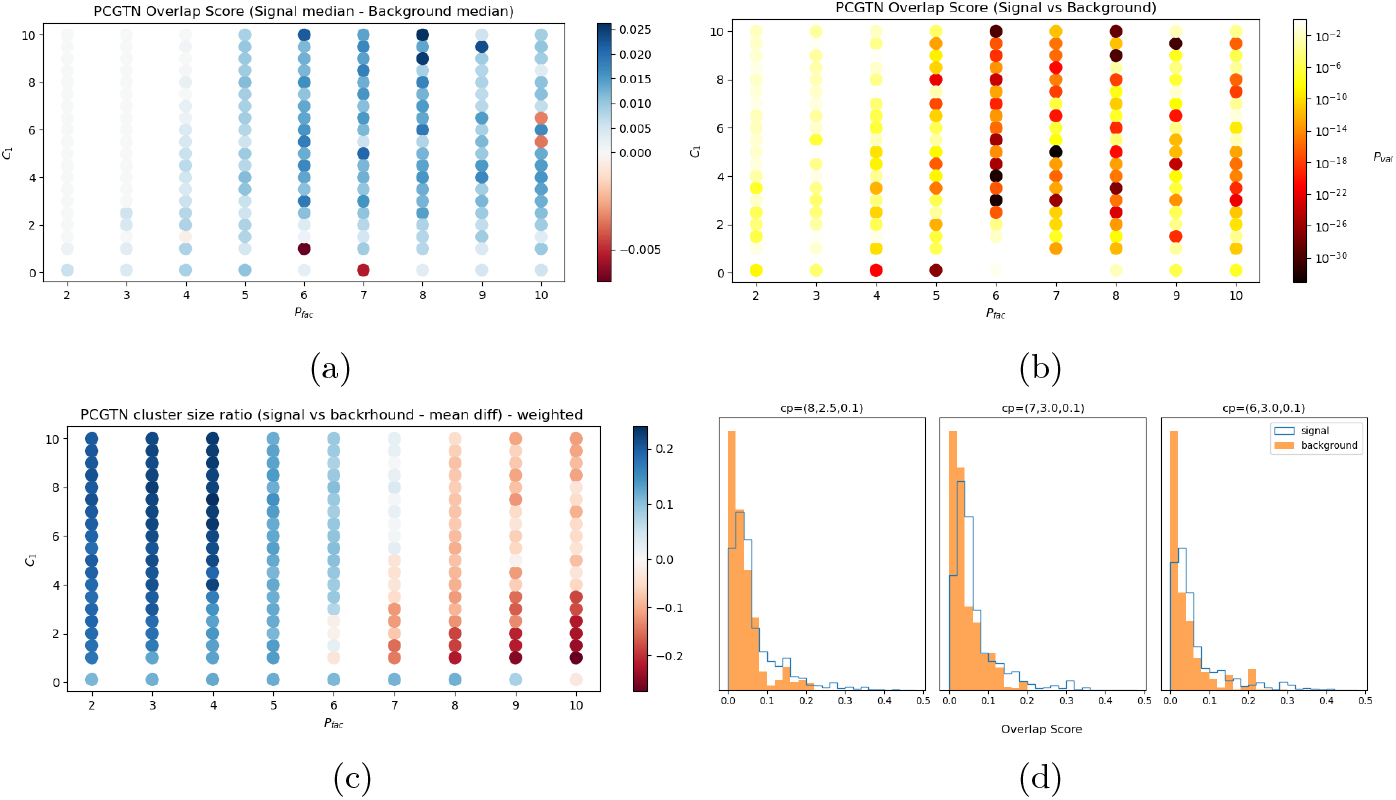
(a) The difference between median overlap score from signal (LLPS) and background proteins. Zeros are in white indicating no difference between medians of two group, positive values are in blue indicating signal group has a larger median, negative values are in red indicating background group has a larger median. (b) Overlap score right hand Mann-Whitney p-value, darker color indicating a smaller p-value as more significant that signal group has a larger overlap score than the background group. (c) The difference between mean value of |1-cluster size ratio | of signal and background proteins. Zeros are in white indicating no difference between two group, positive values are in blue indicating background proteins have cluster size ratio closer to 1, and negative values are in red indicating signal proteins have cluster size ratio closer to 1. (d) For three of the best models from the performance test, we show the distribution of overlap scores for signal and background proteins. See also Figure S3 and S4.

In Figure 5 (a), for the majority of the *cp*-parameters, the median overlap score of signal proteins is larger than that of the background proteins. In (b), the p-value is also significant for almost the entire parameter space. In (c), we observed that for the majority of *cp*− parameters, the sizes of detected clusters is closer to the size of disordered regions for signal proteins than background proteins. In addition, we observe a gradual increase of the difference between the signal and background proteins in (a), a gradual increase of the significance of the hypothesis that the overlap score distribution of signal protein is larger than the background proteins in (b), and an increased difference between how close the cluster sizes and disordered region sizes are in the signal and background proteins in (c) towards the lower right triangular part of the parameter space. This pattern of a higher correspondence between detected clusters and annotated disordered regions, as well as the region densely filled with significant p-values (*p*_fac_ = [6, 8], *C*_1_ ∈ [2, 5], and the region corresponding to the best-performing models around *cp* = (8.0, 2.5, 0.2) as mentioned in Section 3.2) matches the pattern we observed in the performance of our model (see Figure 3).

We also found a higher overlap of annotated polar residue rich regions with our detected clusters in signal proteins compared to background proteins. Further results on the alignment of detected clusters with polar residue rich regions can be found in Supplementary Section S3.3.

In Figure 6, we show an example of the refined graph found by our model compared with disordered region annotations, at an interface between multiple disordered regions (blue shaded in protein illustration) and non-disordered regions (red shaded). The graph plots show the residues and their interactions in the corresponding regions. Residues in disordered regions are shown as large points, while residues in non-disordered regions are shown as small points. Different clusters detected by our model are denoted by various colors, which are kept consistent between the protein illustration and the graph plot. The graph demonstrates that the model accurately detected the central folded structure, grouping it into two clusters marked blue and green. We highlight the interfaces detected by our model with arrows, which match the annotation up to a few residues. Additionally, our model identified another folded structure, marked in orange in the region indicated by the dashed circle. Although this orange cluster contains some residues classified as disordered according to the annotation, our model successfully cut the connection between this subregion and the large disordered region attached to it. We note that there are only a small number of ‘incorrect’ connections, *i* .*e*., edges between a disordered region and a non-disordered region (bright green edges), and these typically involve the same nodes. Across all scaffold proteins, there are only 1.49 *±* 1.58% incorrect edges.

**Figure 6:**
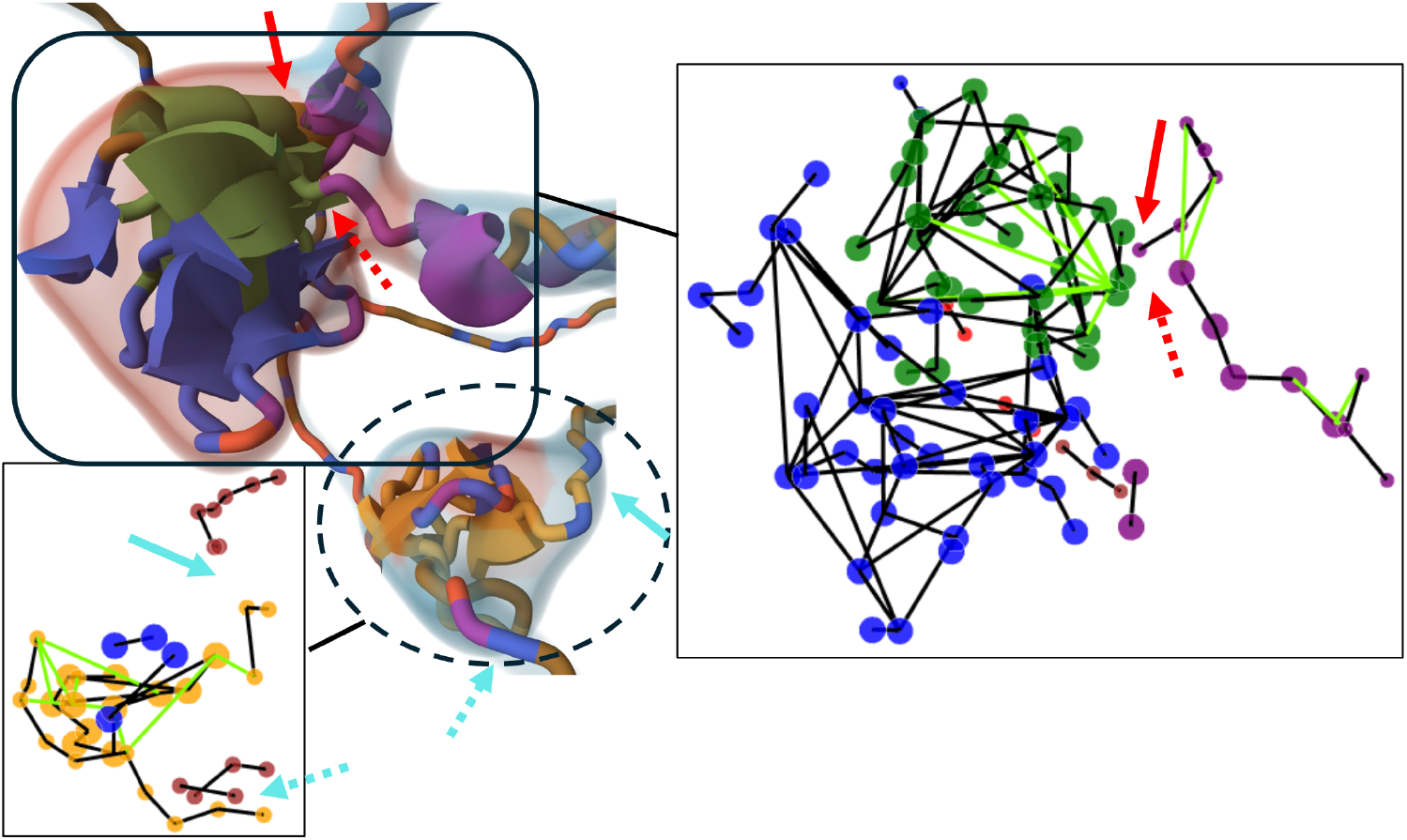
Boundaries of disordered regions in the protein with UniProt ID P35637, as detected by our model (indicated by a lack of edges between different clusters), are compared with the annotations provided in UniProt (represented by shaded colors in the protein illustration and point size in the graphs). The bright green edges highlight incorrect connections, as described in the main text.

## 5 Discussion

In this work, we leveraged a key characteristic of protein LLPS, which is primarily driven by domains such as IDRs, to accurately predict which proteins undergo LLPS. We proposed a GP-GNN framework and demonstrated its effectiveness for LLPS prediction by identifying these crucial domains as subgraphs. To this end, we formulated a similarity measure that captures the probability of nodes pairs jointly exhibiting characteristic patterns within larger systems. We used this measure to remove the connections between nodes that are unlikely to be related for the given task. To enhance the flexibility of our model in adapting to diverse tasks, we leveraged a linear layer to extract meta-features. The usage of meta-features enables the training of the similarity measure with the task labels, thereby facilitating task-specific adjustments in the model’s predictions. We have shown on multiple datasets that our model achieves state-of-the-art performance for protein LLPS prediction, including regulator and scaffold LLPS proteins, with PDB and AlphaFold generated graph-structures. Our approach may also be applied more generally to optimize GNN structure in other domains by identifying task relevant subgraphs, and in other work, we have shown that a similar approach achieves state-of-the-art performance for spatial transcriptomics expression prediction from pathology images, while identifying regions of the tumor and micro-environment that align well with pathologist annotations [50].

For all the datasets considered in this work, adding a clustering layer based on the similarity measure we formulated and a PLL improves the test accuracy by at least 5% compared to conventional graph neural network models. From the model architecture point of view, further investigation of alternative clustering approaches and similarity measures (incorporating, for instance, node interaction or amino acid k-mer terms) may lead to further gains in performance. From the practical training side, increasing the number of samples in the variational optimization, increasing the number of meta-epochs, and additional optimization of network architecture and hyperparameter settings could further improve the performance of the model.

To our best knowledge, our model is the first graph-based predictor for LLPS, demonstrating superior performance compared to existing LLPS predictors in the task of predicting both regulator and scaffold proteins. In addition, we interpreted and validated the clusters identified by our model by aligning them with biological annotated protein disordered regions and polar-rich regions. Notably, the detected clusters successfully match these important biological regions, which may be the principal reason for the improved prediction accuracy. The capability of our model to identify meaningful underlying subgraphs can be further employed to gain additional insights into systems of interest. As an example of downstream analysis, one could align the detected clusters with all annotated protein regions already known to be associated with LLPS proteins. Then, of the remaining novel regions, those with cluster properties, such as 3D protein structure motifs, which are more common in the LLPS proteins could be treated as hypotheses for novel functional regions, and investigated through experimental or computational validation, thereby advancing our understanding of this biological phenomenon.

## Supporting information

folder containing the synthetic dataset and lists of entries of the protein datasets

## Resource Availability

### Data and Code Availability

All original code has been deposited at https://github.com/gersteinlab/GraphPartition_SBGNN. The dataset used for model training can be found in the supplemental information. Any additional information are available from the lead contact upon reasonable request.

## Acknowledgement

We acknowledge support from the NIH and from the AL Williams Professorship funds. We would also like to thank Lawrence H. Phillips for valuable discussions.

## Footnotes

1 The source code is available at https://github.com/gaoyuanwang1976/ProteinGraph_construction.git

2 The small region densely filled with many of the best validation accuracies included *cp* = (9, 3, 0.1), (8, 2.5, 0.1), (8, 2.5, 0.2), (8, 2.5, 0.3), (8, 2.5, 0.4), (8, 3, 0.1), (8, 3.5, 0.1), (7.0, 3.0, 0.1) and (6.0, 2.5, 0.1).

