## supplementary notes for "A Variational Graph Partitioning Approach to Modeling Protein Liquid-liquid Phase Separation"

#### S1 Supplementary information for synthetic dataset

We design a graph classification task with similar characteristics to the protein LLPS task, but where we know the ground-truth functional domains. Particularly, our synthetic data consists of 100 positive and 100 negative graphs each of size 180 nodes. In all graphs, we randomly select 3 foreground clusters, whose sizes are sampled from a normal distribution with mean size 30, and standard deviation 5. Within the same foreground cluster, an edge is assigned between a pair of nodes with probability 0.1, and 0.01 otherwise. We reject any graphs which are not connected. We sample 2D features for each node. For the positive class, the feature is drawn from $\mathcal{N}(0.1 * \mathbf{1}, \mathbf{I})$ for nodes in the foreground clusters and $\mathcal{N}(\mathbf{0}, \mathbf{I})$ otherwise. For the negative class, 90 random nodes have feature from the former distribution and the other 90 nodes from the latter; hence the positive and negative examples have the same global feature statistics, but the 'shifted features' are aligned with the graph clusters only in the positive graphs. We include below results from our GP-GNN framework applied to a synthetic graph classification task.

##### S1.1 Synthetic graph classification results

We compare the basic GNN, CG and PCG models with models trained on fractionized graphs using ground-truth cluster information (TCG), and random dropout GCN with the same average node degree as TCG. The original node degree is 6.15 and the node degree of TCG graphs is 3.8. We train our models for 100 random initializations and report the average and top performance in Table S1. We observe that the GCN trained on TCG clearly outperforms the GCN trained on full graphs and random dropout, confirming that graph sparsification is maximally beneficial when aligned with functionally relevant subgraphs. We further trained the model on fractionized graphs obtained with GP-GNN. Our test covered a wide range of $cp$s and a comprehensive screening was done for $p_{\text{fac}} \in [3, 5]$, $C_1 \in [0.8, 1.6]$ and $C_2 \in [0.7, 1]$. We repeat the training of the CG models 105 times and separate them into 7 groups. For PCG models, we stop the training once $\sigma$ drops below 0.1.

Table S1: Performance of various models on the synthetic dataset. The average and peak performance of basic GCN models, models trained on fractionized graphs with ground-truth (TCGCN), models with edges dropped at random, and models with adaptive clustering / trained pre-linear layers (PCGCN) are compared. The best average performance is highlighted in bold.

| | GCN | TCGCN | GCN random dropout 38% | CGCN | PCGCN |
|---|---|---|---|---|---|
| mean train accu. | $0.633 \pm 0.151$ | $0.807 \pm 0.158$ | $0.74 \pm 0.171$ | $0.889/\pm 0.109$ | $0.877 \pm 0.141$ |
| mean val. accu. | $0.576 \pm 0.021$ | $0.628 \pm 0.036$ | $0.597 \pm 0.043$ | $0.662 \pm 0.032$ | $0.676 \pm 0.037$ |
| mean test accu. | $0.563 \pm 0.064$ | $0.603 \pm 0.046$ | $0.551 \pm 0.066$ | $0.578 \pm 0.063$ | $\mathbf{0.616 \pm 0.052}$ |
| top model test accu. | 0.589 | 0.633 | 0.592 | 0.67 | 0.633 |

The best model is PCGCN with $cp = (4, 1.2, 0.8)$ with an average performance of 0.616±0.052,

Table S2: Average performance (test accuracy) of baseline models on the *Regulator PDB* dataset. The test accuracy is shown for basic GCN and GTN, random dropout GCN and GTN, as well as for the logistic regression model.

| drop % | basic (0%) | 50% | 70% | 90% | logistic regression |
|---|---|---|---|---|---|
| GCN | $0.65 \pm 0.034$ | $0.651 \pm 0.042$ | $0.657 \pm 0.056$ | $0.673 \pm 0.06$ | 0.494 |
| GTN | $0.656 \pm 0.041$ | $0.649 \pm 0.0416$ | $0.641 \pm 0.044$ | $0.646 \pm 0.052$ | |

and a top-model performance of 0.633, which is comparable to the performance when using ground-truth cluster information, confirming that our GP-GNN is able to learn models that perform close to the maximum for this synthetic class of problem.

### S2 Supplementary information for protein LLPS classification

#### S2.1 Protein dataset

The list of the regulator protein dataset with ground truth 3D structure, including their PDB IDs and labels, can be found in `regulator_list_PDB_label.dat`.

The list of the regulator protein dataset with AlphaFold predicted 3D structure, including the proteins' UniProt IDs and labels, can be found in `regulator_list_UniProt_label.dat`.

The list of the scaffold protein dataset with AlphaFold predicted 3D structure, including the proteins' UniProt IDs and labels, can be found in `scaffold_list_UniProt_label.dat`.

We present the chain length distribution of the positive and negative samples in the three datasets in Figure S1, and the amino acid composition in Figure S2. As discussed in the main text, we used a baseline logistic regression model to estimate the impact of biases in these basic protein statistics between the signal and background samples.

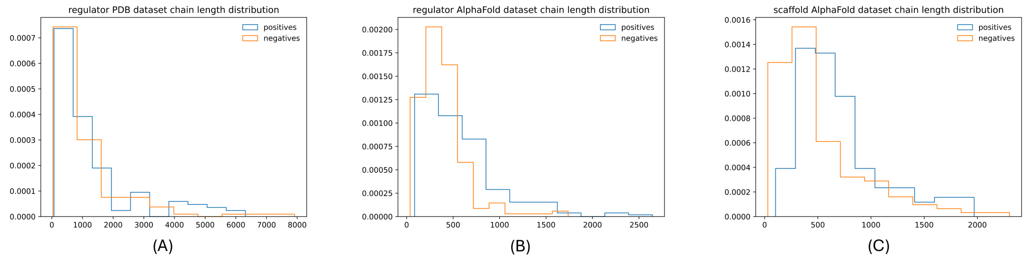


Figure S1: The chain length distribution of the signal proteins is compared with that of the background proteins.

#### S2.2 Additional performance tables

Here, we present the classification accuracy of various models, analogous to the performance tables on AUC in the main text. For *Regulator PDB* dataset, the accuracy of prediction is presented in Table S2 and Table S3. For *Regulator AlphaFold* dataset, it is presented in Table S4. And for *Scaffold AlphaFold* dataset, the classification accuracy on the test dataset can be found in Table S5.

Table S3: *Regulator PDB* dataset, PCGCN and PCGTN models. Average performance of top 3 *cp*-parameters for PCGCN and PCGTN is presented.

| | val. accu. | test accu. | CG test accu |
|---|---|---|---|
| **PCGCN** | | | |
| 2, 0.1, 0.2 | $0.821 \pm 0.036$ | $0.681 \pm 0.042$ | $0.652 \pm 0.052$ |
| 2, 0.1, 0.1 | $0.817 \pm 0.021$ | $0.716 \pm 0.036$ | $0.663 \pm 0.049$ |
| 1, 0.2, 0.1 | $0.813 \pm 0.023$ | $0.7 \pm 0.041$ | $0.681 \pm 0.06$ |
| average | - | $0.699 \pm 0.04$ | $0.666 \pm 0.054$ |
| **PCGTN** | | | |
| 1, 0.1, 0.1 | $0.824 \pm 0.028$ | $0.668 \pm 0.033$ | |
| 1, 0.2, 0.1 | $0.815 \pm 0.022$ | $0.67 \pm 0.024$ | |
| 1, 0.1, 0.4 | $0.79 \pm 0.036$ | $0.672 \pm 0.051$ | |
| average | - | $0.67 \pm 0.038$ | |

Table S4: Performance on the *Regulator AlphaFold* dataset for various models. Average performance of the three *cp*-parameters with the highest validation performance for PCGCN and PCGTN is presented.

| **PCGCN** | | | **PCGTN** | | |
|---|---|---|---|---|---|
| *cp* | val. accu. | test accu. | *cp* | val. accu. | test accu. |
| 8, 2.5, 0.1 | $0.722 \pm 0.027$ | $0.675 \pm 0.045$ | 7, 3.0, 0.1 | $0.737 \pm 0.028$ | $0.693 \pm 0.032$ |
| 7, 3.0, 0.1 | $0.718 \pm 0.018$ | $0.677 \pm 0.042$ | 8, 2.5, 0.2 | $0.736 \pm 0.028$ | $0.695 \pm 0.044$ |
| 6, 2.5, 0.1 | $0.714 \pm 0.029$ | $0.674 \pm 0.03$ | 8, 3.5, 0.1 | $0.734 \pm 0.044$ | $0.726 \pm 0.022$ |
| average | - | $0.675 \pm 0.04$ | average | - | $0.705 \pm 0.034$ |
| | basic GCN | basic GTN | logistic regression | | |
| test accu. | $0.656 \pm 0.036$ | $0.671 \pm 0.03$ | 0.587 | | |

Table S5: The test accuracy of various models on the *Scaffold AlphaFold* dataset is presented.

| | | | |
|---|---|---|---|
| basic GCN | basic GTN | random dropout GTN (34%) | logistic regression |
| $0.726 \pm 0.054$ | $0.76 \pm 0.052$ | $0.754 \pm 0.051$ | 0.659 |
| PCGCN(7,3.0,0.1) | CGTN(8,2.5,0.2) | PCGTN(8,2.5,0.2) average | PCGTN(8,2.5,0.2) peak |
| $0.746 \pm 0.046$ | $0.696 \pm 0.057$ | $0.811 \pm 0.035$ | 0.841 |

#### S2.3 Additional metrics for selected models on scaffold dataset

For our peak-performing model, PCGTN (8, 2.5, 0.2), we provide additional prediction performance metrics, including Matthew's correlation coefficient (MCC), precision, recall, specificity and F1 score, on the *Scaffold AlphaFold* dataset in Table S6. The definition of the performance metrics can be found in Equation S1, where TP, TN, FP, and FN represent True Positive, True Negative, False Positive, and False Negative, respectively.

Table S6: Prediction performance of PCGTN on the *Scaffold AlphaFold* dataset compared with previous sequence-based predictors.

| | PCGTN | FuzDrop [1] | catGRANULE [2] | ParSe [3] |
|---|---|---|---|---|
| AUC | 0.92 | 0.886 | 0.753 | 0.751 |
| Precision | 0.85 | 0.805 | 0.659 | 0.707 |
| Recall | 0.829 | 0.805 | 0.659 | 0.707 |
| Specificity | 0.854 | 0.805 | 0.659 | 0.707 |
| F1 score | 0.839 | 0.805 | 0.659 | 0.707 |
| MCC | 0.683 | 0.61 | 0.317 | 0.415 |

$$\begin{aligned}
\text{Specificity} &= \frac{\text{TN}}{\text{TN}+\text{FP}} \\
\text{Precision} &= \frac{\text{TP}}{\text{TP}+\text{FP}} \\
\text{Recall} &= \frac{\text{TP}}{\text{TP}+\text{FN}} \\
\text{F1 score} &= 2\times\frac{\text{Precision}\times\text{Recall}}{\text{Precision+Recall}} \\
\text{MCC} &= \frac{\text{TP}\times\text{TN}-\text{FP}\times\text{FN}}{\sqrt{(\text{TP}+\text{FP})\times(\text{TP}+\text{FN})\times(\text{TN}+\text{FP})\times(\text{TN}+\text{FN})}}
\end{aligned} \tag{S1}$$

#### S2.4 Comparison with prior LLPS results

As described in the main text (Section 3.3 **Comparison to previous predictors**), we compared our model with existing methods for phase separation protein (PSP) prediction [4, 5, 6, 7]. To this end, we assessed likelihood scores of the PSPs and background proteins in our test datasets being classified as PSP with our model. We conducted this analysis separately for both the regulator proteins and scaffold proteins using the AlphaFold predicted structure information. The score we used to rank the proteins is the class one (signal) probability of the two class probability outputs (we used the argmax function to obtain the final predicted output from the two class probability output).

We applied the optimal PCG models specific to the *Regulator AlphaFold* dataset and *Scaffold AlphaFold* dataset correspondingly. We obtained the results separately for the 15 samples, and ranked the score within each sample. If the second background protein is found to have rank $n$, then there are $n-2$ PSPs ranked before the second background protein. The recall is then $(n-2)/N_s$, where $N_s$ is the number of total PSPs. The mean and standard deviation of the recalls from the 15 samples were provided in the main text.

As a further comparison, we bench-marked our performance directly against several prior methods applied to the test partition for our scaffold dataset. The results for this comparison are shown on Table S6. For the three methods compared (FuzDrop [1], catGRANULE [2] and ParSE [3]), web interfaces were provided (we were unable to access the web interface for PSPredictor, since the address was listed as not secure). The results shown are calculated using the ranking of proteins in our test set provided by each method, using the test set sequences as inputs (for metrics other than AUC, a median threshold is chosen to ensure equal numbers of predicted positives and negatives, since our test set is equally balanced). Finally, we note that a more extensive set of eleven recent LLPS prediction methods was compared in [8], where the results given on their *P_All vs N_Human* dataset may be regarded as comparable to our scaffold dataset, since the positives are known scaffold proteins (not limited to the human proteome), while the negatives are sampled uniformly from PDB without restriction on the degree of resolution (although the negatives in [8] are restricted to the human proteome, while ours are not; we note, however, that our dataset may be harder for this reason, since the statistics of the positive and negative examples are expected to be more closely matched). The following methods are reported to achieve best AUC scores on the *P_All vs N_Human* dataset: LLPhyscore ($0.88 \pm 0.005$), PSPredictor ($0.83 \pm 0.009$), PSPer ($0.82 \pm 0.012$), FuzDrop ($0.79 \pm 0.007$), and catGRANULE ($0.79 \pm 0.008$). We note that these scores are comparable to the distribution of scores achieved across methods on our dataset in Table Table S6, and that the performance of the PCGTN (0.92 AUC) is competitive in the context of the latter.

### S3 Supplementary information for validation against annotated protein regions

#### S3.1 Disordered region annotation validation results for $C_2$ not equal to $0.1$

Additional screening for $C_2 \in \{0.2, 0.3, 0.4, 0.5, 1.0, 2.0, 3.0, 5.0\}$ has been conducted for the top performing models that have ($p_{\text{fac}}$,$C_1$)$\in$ [$(6, 1.0)$, $(8, 2.5)$]. The results for two metrics are presented in Table S7: the right-hand Mann-Whitney p-value for the hypothesis that signal proteins have higher overlap scores than background proteins, and the median difference in overlap score between signal and background proteins. For nearly all *cp*-parameters shown here, the signal proteins have markedly larger distribution of overlap scores compared to background proteins.

Table S7: Disordered region annotation validation results for $C_2 \neq 0.1$. For all *cp*-parameters, signal proteins have higher overlap scores compared to background proteins, and we observed significant right-hand Mann-Whitney p-value for 15 out of 16 *cp*-parameters.

| *cp* | right-hand Mann-Whitney p-val. | median diff. |
|---|---|---|
| 6,1.0,0.2 | $8.279008 \times 10^{-15}$ | 0.026141 |
| 6,1.0,0.3 | $6.074975e \times 10^{-12}$ | 0.027557 |
| 6,1.0,0.4 | $2.078727 \times 10^{-13}$ | 0.027558 |
| 6,1.0,0.5 | $1.440727 \times 10^{-12}$ | 0.027473 |
| 6,1.0,1.0 | $7.275033 \times 10^{-09}$ | 0.025974 |
| 6,1.0,2.0 | $5.411554 \times 10^{-07}$ | 0.017136 |
| 6,1.0,3.0 | $9.896033 \times 10^{-11}$ | 0.018196 |
| 6,1.0,5.0 | $3.672743 \times 10^{-10}$ | 0.027224 |
| 8,2.5,0.2 | $1.110580 \times 10^{-01}$ | 0.013221 |
| 8,2.5,0.3 | $1.661899 \times 10^{-03}$ | 0.022368 |
| 8,2.5,0.4 | $6.924865 \times 10^{-06}$ | 0.014545 |
| 8,2.5,0.5 | $1.296786 \times 10^{-16}$ | 0.031746 |
| 8,2.5,1.0 | $5.532527 \times 10^{-07}$ | 0.026915 |
| 8,2.5,2.0 | $5.790740 \times 10^{-09}$ | 0.030433 |
| 8,2.5,3.0 | $1.476415 \times 10^{-09}$ | 0.028867 |
| 8,2.5,5.0 | $5.779626 \times 10^{-09}$ | 0.030970 |

#### S3.2 Alternative overlap score and cluster size error method and results

As outlined in Section 4 **Validation against annotated protein regions** of the main text, our model's identified clusters are validated using the weighted Jaccard Index and a weighted analysis of cluster size ratio error. In addition, we present further results using a modified approach for both the Jaccard Index and cluster size ratio error. For a given protein, rather than computing a weighted average, we select the highest Jaccard Index from each disordered region and then choose the maximum value from these. This method focuses on identifying the most accurately overlapping region among all clusters. Similarly, we apply this modification to the calculation of cluster size ratio error. Here, instead of averaging, we select the smallest cluster size ratio error from each disordered region and then choose the smallest value from among these, aiming to pinpoint the most precise cluster size approximation across all regions. We show the results of this additional analysis in Figure S3.

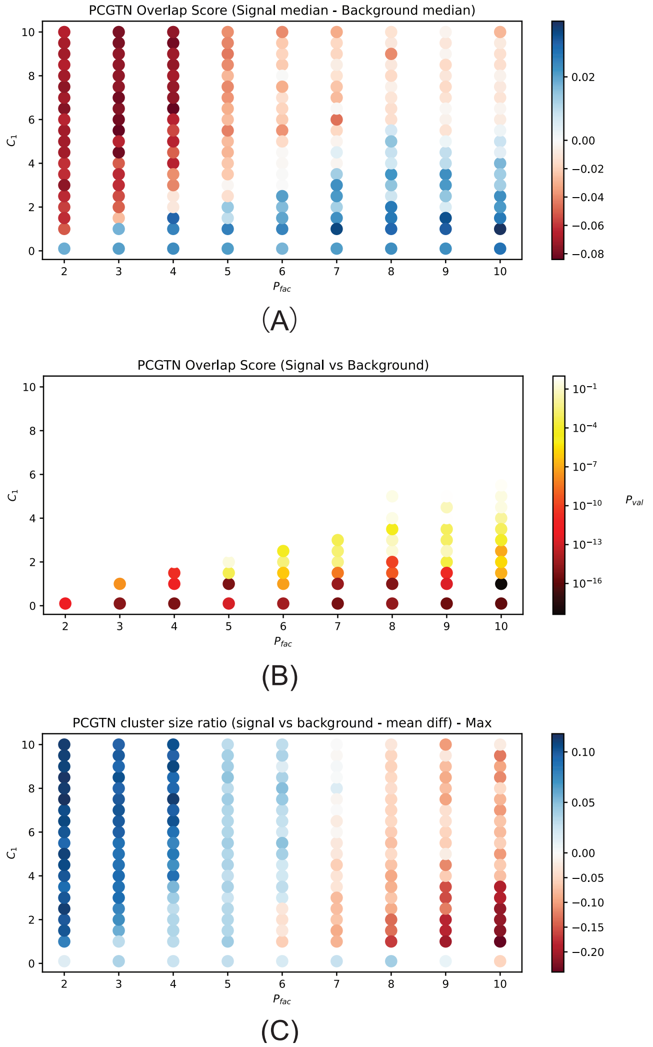


Figure S3: (A) The difference between median overlap score from signal (LLPS) and background proteins. Zeros are in white indicating no difference between medians of two group. (B) Overlap score right hand Mann-Whitney p-value, darker color indicating a smaller p-value as more significant that signal group has a larger overlap score than the background group. (C) The difference between mean value of |1- cluster size ratio| of signal and background proteins. Zeros are in white indicating no difference between two group, positive values are in blue indicating background proteins have cluster size ratio closer to 1, and negative values are in red indicating signal proteins have cluster size ratio closer to 1.

#### S3.3 Overlap with polar residue rich regions

We further assess the biological insights obtained by our model by comparing the clusters detected against polar residue rich regions, as annotated in UniProt. We calculated the overlap score between polar residue rich regions and clusters detected by PCGTN using the same approach described in the main text (see Section 4 in the main text, "Overlap assessment using Jaccard Index").

In Figure S4 (A), one can see that for the majority of the *cp*-parameters, the median overlap score of signal proteins is larger than that of the background proteins. In (B), the right-hand Mann-Whitney p-value for the hypothesis that signal proteins have higher overlap with polar residue rich regions than background protein is also significant for almost the entire parameter space.

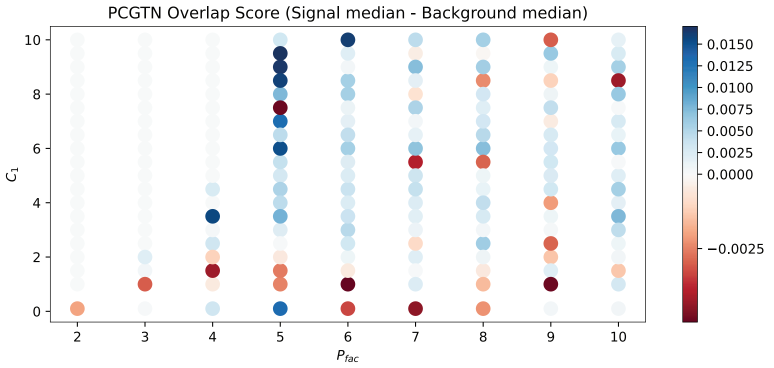


(A)

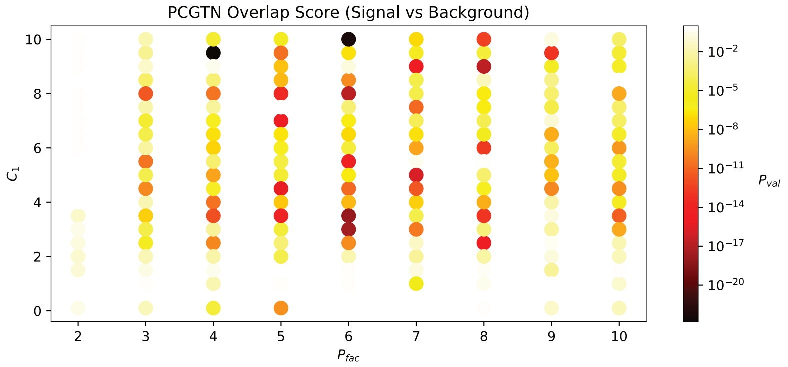


(B)

Figure S4: (A) The difference between median overlap scores from signal and background proteins. Zeros are in white indicating no difference between medians of two groups, positive values are in blue indicating signal group has a larger median, and negative values are in red indicating background group has a larger median. (B) Overlap score right hand Mann-Whitney p-values, with darker colors indicating a smaller p-value (more significant), for the signal group having a larger overlap score than the background group.

### S4 Code availability

In addition to the interactions considered by the proteingraph package[9], we included also planar pi-pi interaction as described in [4]. The code for protein graphs generation is available at `https://github.com/gersteinlab/ProteinGraph_construction`.

To prepare protein graphs with AlphaFold predicted structure information and store them in `/graph_test`, run

```
mkdir graph_test
python GNN_createGraph.py -d scaffold_list_UniProt_label.dat --graph_path \
graph_test/ --pdb_path scaffold_AlphaFold_pdb/ -i AlphaFold
```

The codes to run our GP-GNN models, the basic GNN, and random dropout GNN are available at `https://github.com/gersteinlab/GraphPartition_SBGNN`. The additional files from this study to run the code for LLPS proteins (with AlphaFold predicted 3D structure) or the synthetic dataset can be found in the supplement material as well.

- Run basic GNN on the scaffolds
  ```
  python PlainGNN.py -d scaffold_list_UniProt_label.dat --graph_path \
  scaffold_AlphaFold/
  ```
- Run random dropout GNN
  ```
  python RandomDropGNN.py -d scaffold_list_UniProt_label.dat \
  --graph_path scaffold_AlphaFold/
  ```
- Run PCGCN
  ```
  python SBGNN_ray.py -d scaffold_list_UniProt_label.dat --graph_path \
  scaffold_AlphaFold/
  ```
- For the synthetic dataset, use the following command line arguments
  ```
  -d synthetic/graph_list.dat --graph_path synthetic/graph/
  ```

[9] https://github.com/ericmjl/protein-interaction-network/. Version 0.3.0.

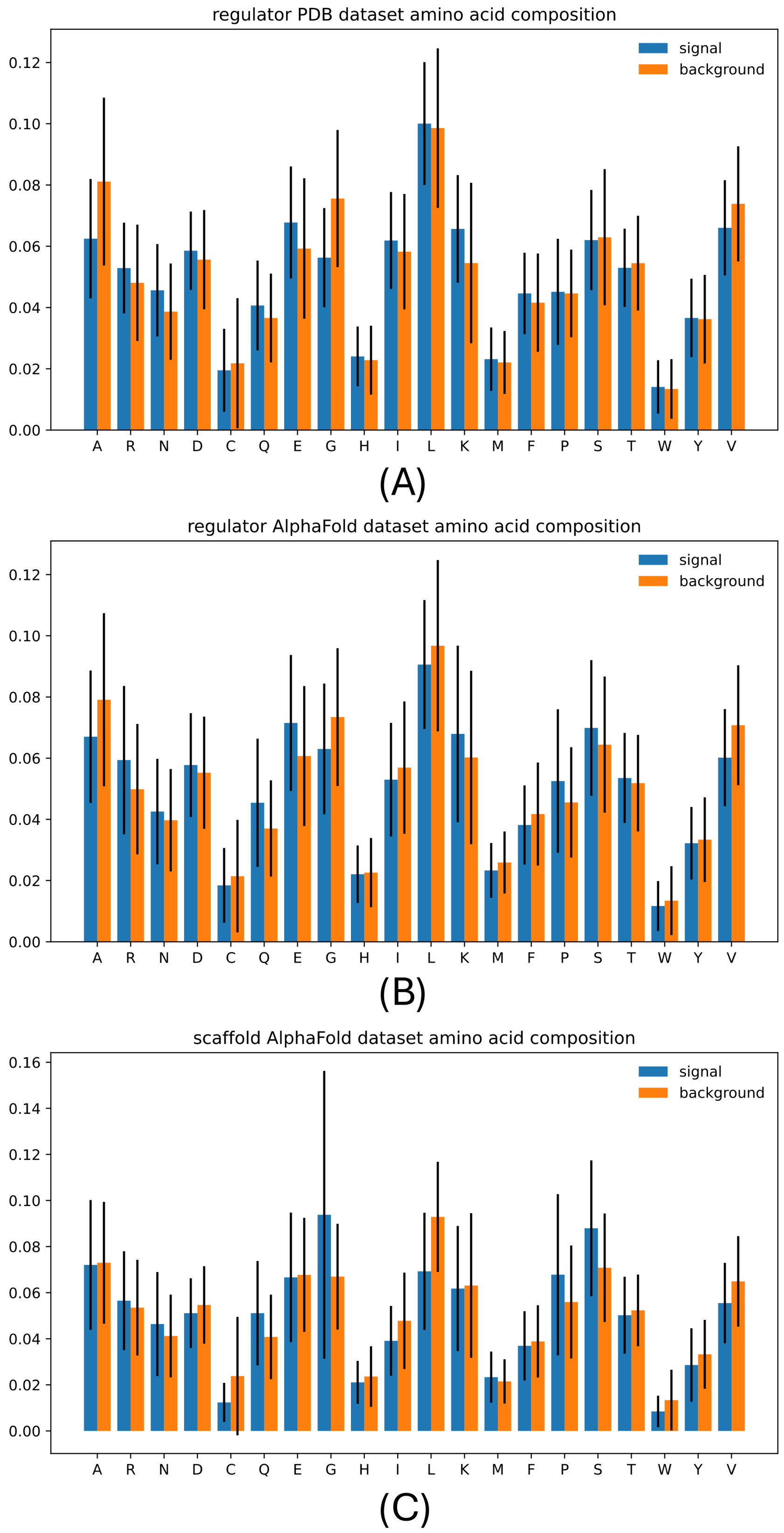


Figure S2: The amino acid composition of the signal proteins is compared with that of the background proteins.
